# Small Intestinal Goblet Cells Control Humoral Immune Responses and Mobilization During Enteric Infection

**DOI:** 10.1101/2024.01.06.573891

**Authors:** Devesha H. Kulkarni, Khushi Talati, Elisabeth L. Joyce, Hrishi Kousik, Dalia L. Harris, Alexandria N. Floyd, Vitaly Vavrinyuk, Bibianna Barrios, Sreeram Udayan, Keely McDonald, Vini John, Chyi-Song Hsieh, Rodney D. Newberry

**Author notes:** Corresponding Authors: Devesha H. Kulkarni, PhD, Rodney D. Newberry, MD.

## Abstract

Humoral immune responses within the gut play diverse roles including pathogen clearance during enteric infections, maintaining tolerance, and facilitating the assemblage and stability of the gut microbiota. How these humoral immune responses are initiated and contribute to these processes are well studied. However, the signals promoting the expansion of these responses and their rapid mobilization to the gut mucosa are less well understood. Intestinal goblet cells form goblet cell-associated antigen passages (GAPs) to deliver luminal antigens to the underlying immune system and facilitate tolerance. GAPs are rapidly inhibited during enteric infection to prevent inflammatory responses to innocuous luminal antigens. Here we interrogate GAP inhibition as a key physiological response required for effective humoral immunity. Independent of infection, GAP inhibition resulted in enrichment of transcripts representing B cell recruitment, expansion, and differentiation into plasma cells in the small intestine (SI), which were confirmed by flow cytometry and ELISpot assays. Further we observed an expansion of isolated lymphoid follicles within the SI, as well as expansion of plasma cells in the bone marrow upon GAP inhibition. S1PR1-induced blockade of leukocyte trafficking during GAP inhibition resulted in a blunting of SI plasma cell expansion, suggesting that mobilization of plasma cells from the bone marrow contributes to their expansion in the gut. However, luminal IgA secretion was only observed in the presence of *S. typhimurium* infection, suggesting that although GAP inhibition mobilizes a mucosal humoral immune response, a second signal is required for full effector function. Overriding GAP inhibition during enteric infection abrogated the expansion of laminar propria IgA+ plasma cells. We conclude that GAP inhibition is a required physiological response for efficiently mobilizing mucosal humoral immunity in response to enteric infection.

## INTRODUCTION

A single layer of columnar epithelial cells separates the gut luminal contents, containing trillions of microbes, their products, and diet from the largest collection of immune cells in the body. In the steady state, the gut immune system, comprised in part of the intestinal epithelium and underlying immune system, promotes tolerance in this potentially harmful environment, which is in part driven by the presence of cellular populations with tolerogenic properties, the production of tolerogenic cytokines, and the presence of microbial and dietary substances promoting tolerance [1–3]. Conversely, when faced with a pathogen, the gut immune system rapidly mounts a protective immune response characterized by T effector responses and humoral immunity, which is believed to largely be driven by pathogen recognition. While great progress has been made in dissecting the role of specific immune cell subsets, cytokines, and other factors promoting tolerance or effector responses, how the gut immune system rapidly overcomes this strong tolerogenic environment to mount an inflammatory response and how this inflammatory response resolves to return to tolerance are not well understood.

IgA production and secretion at mucosal surfaces constitutes one of the largest immunoglobulin pools in the body [4, 5]. Gut luminal IgA has steady state functions of helping to maintain the gut barrier, sequestering gut microbes from the epithelium, providing niches for beneficial gut microbes, and promoting the assembly and structure of the gut microbiota [6, 7]. Further during infection, IgA can be rapidly mobilized to aid in protection from barrier breach and pathogen clearance [8, 9]. IgA can have multiple origins, including being derived from classical B2 B cells with CD4+ T cell help within organized lymphoid tissue, derived from B-1 B cells with or without T cell help, and derived from B cells within isolated lymphoid follicles (ILFs) independent of T cell help [10]. The differentiation of B cells into IgA antibody-secreting cells (ASCs) mainly takes place in the gut-associated lymphoid tissue, which include the Peyer’s patches and ILFs of the lamina propria [11]. After differentiation, these IgA-producing ASCs, which includes proliferating plasmablasts and terminally differentiated non-mitotic plasma cells reside in the lamina propria (LP), where they produce IgA to be secreted into the gut lumen and/or contribute to the systemic IgA pool [5, 12].

While it is known that numerous IgA- (and to a lesser extent IgG-) producing ASCs are found in the intestinal mucosa, and that plasma cell and Ig responses are rapidly expanded and mobilized in response to challenges in the gut [13], the factors controlling the gut plasma cell population and coordinating these responses are not well understood. Here, we demonstrate that blocking the ability of goblet cells to form GAPs, a pathway supporting tolerance during homeostasis, promotes the rapid induction of humoral immune responses within the intestine and bone marrow even in the absence of enteric infection. However, in contrast to events during enteric infection, the mobilization of humoral immune responses by inhibition of GAPs does not increase luminal or systemic IgA. Intriguingly, the mobilization of IgA responses during enteric infection was abrogated when GAP inhibition was overridden, indicating that the inhibition of GAPs during enteric infection is part of a physiological effector response central to mobilizing humoral immunity at this mucosal surface during enteric infection.

## RESULTS

### Inhibition of intestinal goblet cell-associated antigen passages induces IgA responses in the gut

Intestinal goblet cells play key roles in intestinal immunity through the secretion of mucus and antimicrobial peptides to form one of the first barriers in defense of the gut epithelium. In addition to these roles, goblet cells can form goblet cell associated antigen passages (GAPs), which function to deliver luminal antigens and goblet cell products to the underlying immune system in the LP, imprint LP antigen-presenting cells (APCs) with tolerogenic capacities, maintain LP T regulatory cells (Tregs), and induce antigen-specific Tregs in the draining lymph nodes [14, 15]. GAPs form by an atypical fluid phase endocytic process delivering luminal contents to the transcytotic pathway [16]. GAP formation is induced by acetylcholine (ACh) acting on the muscarinic ACh receptor 4 (mAChR4) on goblet cells and is inhibited by the activation of the epidermal growth factor receptor (EGFR) on goblet cells, which can occur directly via EGFR ligands or indirectly by transactivation via MyD88 within goblet cells [17]. We have previously reported that during enteric infection, GAPs are inhibited by IL-1β acting on the IL-1R resulting EGFR transactivation via MyD88 [18]. GAP inhibition during enteric infection results in two salient outcomes; 1) preventing the development of inflammatory responses to dietary antigens due to the lack of luminal antigen delivery to the immune system, and 2) limiting the dissemination of the pathogen, which used GAPs as a portal of entry [18].

To evaluate if inhibition of GAPs independent of enteric infection had effects on the gut immune system, we conducted bulk RNA-sequencing of the small intestine of mice where GAPs were inhibited by the inducible genetic deletion of mAChR4 on goblet cells [15] (mAChR4^f/f^Math1^Cre*PR^ mice) and Cre-littermate control mice following treatment with RU486 for seven days. Previous studies have demonstrated that the deletion of mAChR4 on goblet cells significantly reduces GAPs but does not affect goblet cell number or mucus thickness [15]. Gene set enrichment analysis of differentially expressed genes by RNA-seq revealed a strong enrichment for genes associated with immune responses with GAP inhibition (**Fig. 1A**). Somewhat surprisingly, as B cell responses were not previously noted to be associated with GAPs, GAP inhibition revealed a strong signal associated with humoral immune responses, including the upregulation of genes such as *PRDM1, IGLC1, CXCL13* (**Fig. 1B**). ELISpot assays revealed a significant increase in SI-LP IgA-producing plasma cells when GAPs were inhibited by inducible deletion of mAChR4 on goblet cells (**Fig. 1C**), when goblet cells and GAPs were deleted by inducible deletion of Math1 on intestinal epithelial cells (**Fig. 1D**), and when GAPs were inhibited pharmacologically by gavage with EGF (**Fig. 1E**) indicating that the induction of humoral immune responses was common to multiple approaches to inhibit/delete GAPs. Expansion of IgA+ plasma cells in the SI-LP of mice with inhibition of GAPs was associated with increased of IgA in LP cells culture *ex vivo* for three days **(Fig. 1F)**, and with increased expression of PIGR by intestinal epithelial cells (**Fig. 1G).** However, while inhibition of GAPs result in modest increase in levels of SI-luminal IgA, the concentration of fecal or serum IgA remained unaltered (**Ext Fig. 1A-C).** In contrast, deletion of goblet cells, which is anticipated to have effects beyond the loss of GAPs, resulted in an increase in luminal and serum IgA, but not serum IgM or IgG **(Ext Fig. 1E-F)**. Consistent with the lack of alterations in luminal IgA when GAPs were inhibited, 16S rRNA sequencing of the small intestinal luminal contents from mAChR4^f/f^Math1^Cre*PR^ and littermate mice controls revealed minimal changes to small intestinal microbial composition at seven days following GAPs inhibition **(Ext Fig. 2)**. Thus, inhibition of GAPs resulted in an unanticipated mobilization of humoral immune responses to the gut.

**Figure 1:**
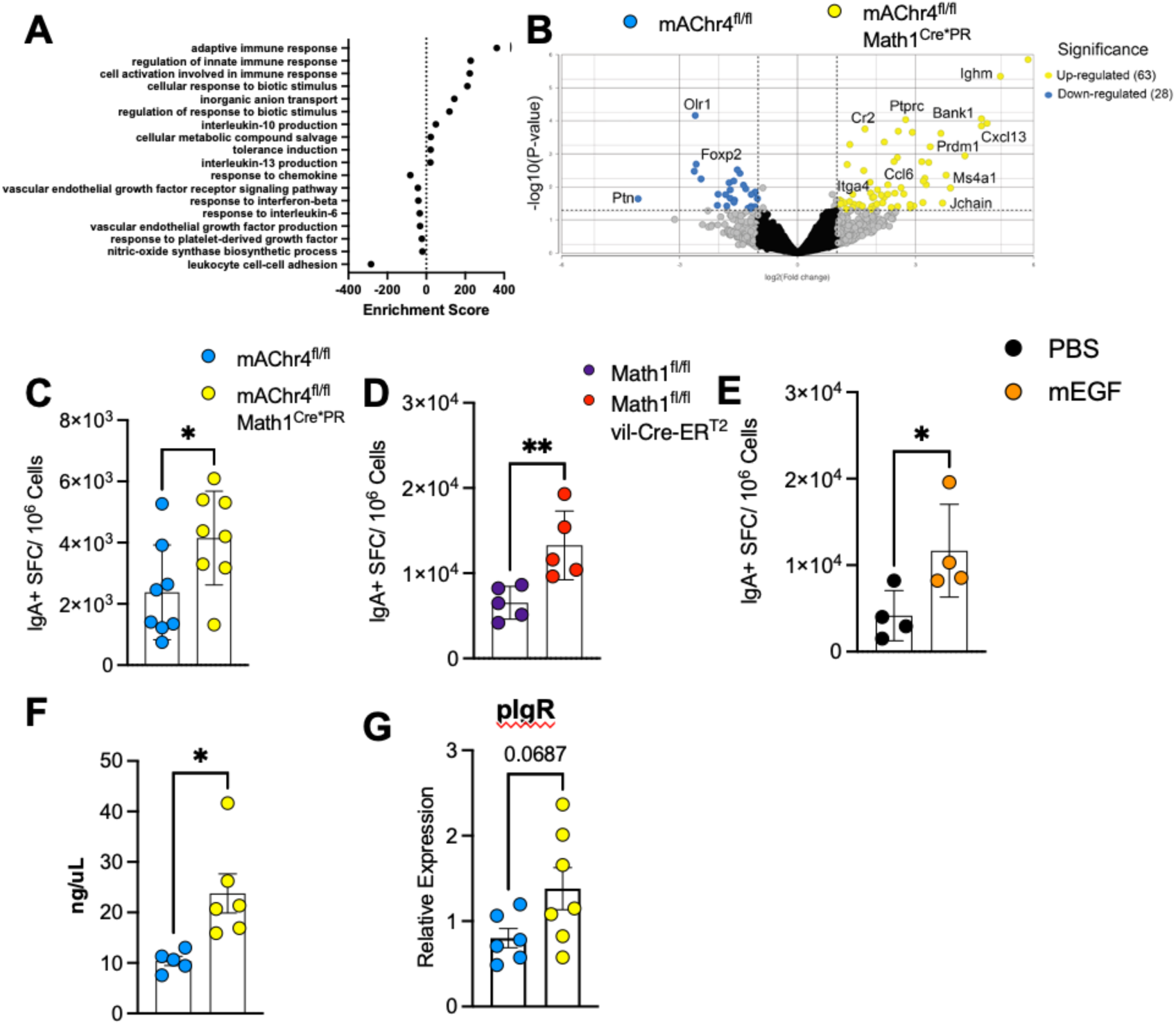
Loss of intestinal goblet cells or inhibition of GAPs results in expansion of plasma cells in small intestinal lamina propria. **(A-B)** Bulk RNAseq was performed on RNA isolated from small intestinal tissue from mAChr4^f/f^ and mAChr4^f/f^ Math1^Cre*PR^ mice treated with RU486 to activate Cre for seven days. (**A**) Graph shows cumulative gene enrichment score upregulated in mAChr4^f/f^ Math1^Cre*PR^ mice (positive enrichment score) compared with mAChr4f/f mice (negative enrichment score). (**B**) Volcano plot depicting DEGs. Yellow dots represent significantly upregulated genes in mAChr4^f/f^ Math1^Cre*PR^ mice and blue dots represent significantly upregulated genes in mAChr4f/f mice. (**C**) Graph depicts small intestinal IgA-secreting cells measured by ELISpot assay in mAChr4f^/f^ and mAChr4^f/f^ Math1^Cre*PR^ mice. (**D**) Small intestinal IgA-secreting cells measured by ELISpot assay in Math1^f/f^ and Math1^f/f^ Vil-Cre-ER^T2^ mice treated with tamoxifen to induce goblet cell deletion. (**E**) Small intestinal IgA-secreting cells measured by ELISpot assay in C57BL/6 mice gavaged with PBS or recombinant EGF for seven days. (**F**) ELISA analysis of IgA concentration in small intestinal lamina propria culture for 3 days. (**G**) Graph depicts relative expression of PIGR in small intestinal tissue. (H) Concentration of BAFF was determined in tissue homogenate. (**C-G**) Data pooled from 2-3 independent experiments. Statistics were calculated by unpaired *t*-test. *p< 0.05, **p<0.01.

**Figure 2:**
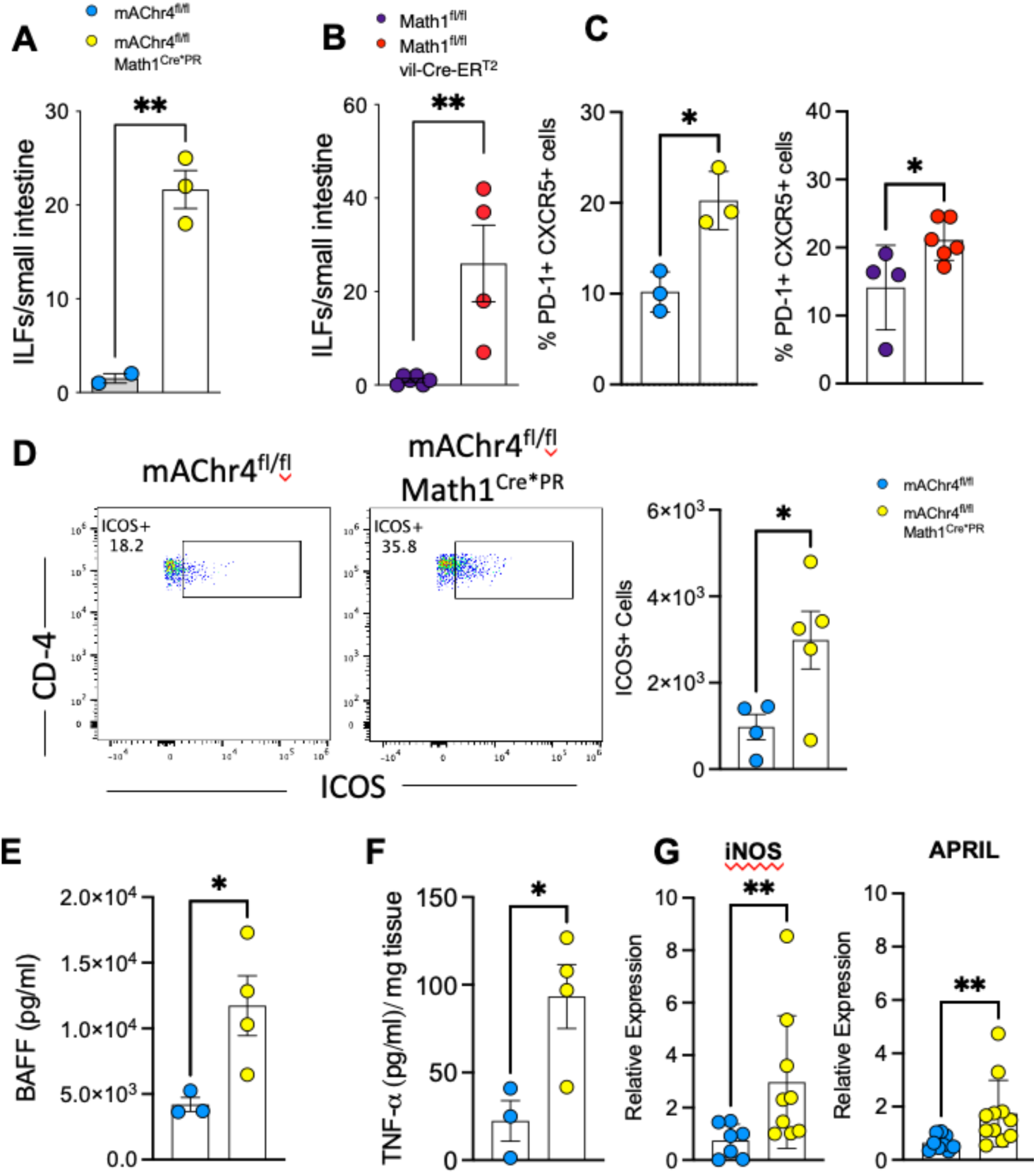
Expansion of Isolated lymphoid follicles and T-helper follicular cells (Tfh) following inhibition of GAPs. **(A)** Densities of small intestinal Isolated Lymphoid Follicles (ILFs) were measured in mAChr4^f/f^ and mAChr4^f/f^ Math1^Cre*PR^ mice and (**B**) Math1^f/f^ and Math1^f/f^ Vil-Cre-ER^T2^ mice. (**C**) Peyer’s Patches were isolated and Tfh cell frequency was determined using flow cytometric analysis based on PD1+ CXCR5+ cells among CD45+ CD3+ CD4+ cells. (**D**) Flow plots show ICOS expression on non-Tfh cells in the small intestinal lamina propria from mAChr4^f/f^ and mAChr4^f/f^ Math1^Cre*PR^ mice. Bar graph adjacent shows quantification of absolute ICOS+ non-Tfh cells. (**E**) Graph depicts quantification of cytokine levels in small intestinal tissue of mAChr4^f/f^ and mAChr4^f/f^ Math1^Cre*PR^ mice. (**F**) Quantitative real time PCR for expression of iNOS and APRIL in RNA isolated from ileal tissue. Data pooled from 2 independent experiments. Statistics were calculated by unpaired *t*-test. *p<0.05, **p<0.01.

### Goblet cells and GAPs orchestrate ILFs and TFH cells in Peyer’s patches

IgM+ naïve B cells migrate from the bone marrow to secondary lymphoid tissues such as Peyer’s patches (PP), where they undergo class switch recombination, express surface Igs of various classes, and progress to become Ig-producing plasma cells [19, 20]. Unlike PP, which develop *in utero* and remain fixed in number throughout life, ILFs develop after birth arising from cryptopatches in response to stimuli and are dynamic, forming, regressing and changing in numbers throughout life [21–24]. B cells within ILFs have been shown to differentiate into IgA-producing plasma cells [11, 21]. Cryptopatches and ILFs are extremes of the continuum of solitary intestinal lymphoid tissues with cryptopatches (∼900/intestine) being the precursors to ILFs (∼100 B220+ clusters; immature ILFs, <5 follicle with dome like structures; mature ILFs) in SPF-housed C57BL/6 mice [21–23]. Only mature ILFs have been observed to give rise to IgA-producing cells [11], and CXCL13 has been found to play a role in the development of ILFs [23]. Given that CXCL13 was significantly upregulated in the non-PP containing small intestinal tissue when GAPs were inhibited (**Fig. 1B**), we evaluated if GAP inhibition induced the development of mature ILFs. In line with the observation that GAP inhibition mobilized an IgA humoral immune response in the gut, we saw that the deletion of goblet cells or the loss of GAPs results in expansion of ILFs in the small intestine (**Fig. 2A-B**). Within the PP, the naïve B cells interact with antigen-primed CD4+ follicular helper T cells (Tfh) [25]. Relatedly we observed that the loss of goblet cells or the inhibition of GAPs results in expansion of Tfh (PD1+ CXCR5+ CD4 T cells) in the PP (**Fig. 2C**). In addition to Tfh, the process of class switch recombination has been reported to occur in extra-follicular regions of the lymph and the spleen, where ICOS-expressing non-Tfh cells are known to promote this process [26, 27]. We therefore assessed ICOS expression on non-Tfh CD4+ T cells and observed an increase in ICOS-expressing CD4+ non-Tfh cells in mice where GAPs were inhibited (**Fig. 2D**). B cell activation factor of the TNF family (BAFF) is a potent B cell survival factor, produced by multiple immune cell types, and BAFF has been shown to be critical for IgA production [28]. We observed that mAChR4^f/f^Math1^Cre*PR^ mice had higher BAFF expression compared to littermates (**Fig 2E**). TNF-α, along with inducible nitric oxide synthase (iNOS) has been shown to regulate IgA class switching [20]. Indeed, we observed that inhibition of GAPs increased TNF-α levels in the gut (**Fig. 2F**). Additionally, the expression of iNOS and APRIL were significantly elevated in mAChR4^f/f^Math1^Cre*PR^ mice compared to littermates (**Fig. 2G**). These data indicate that inhibition of GAPs results in the induction of a generalized program supporting IgA production within the small intestinal mucosal lymphoid tissues and LP.

### Inhibition of GAPs mobilizes B cell precursor populations in the bone marrow

The bone marrow is the site of B cell generation, and recent evidence suggests that B cell maturation can also occur in the bone marrow [29]. B cells have been shown to rapidly accumulate in the bone marrow during acute systemic inflammation [30]. Additionally, a substantial population of mucosal ASCs are found in the bone marrow, where long-lived ASCs reside [31, 32]. Further, studies have demonstrated the homing of gut ASCs to the bone marrow following oral immunization [33]. Therefore, we analyzed the subsets within the B cell development pathway in the bone marrow when GAPs are inhibited to determine if this humoral immune response mobilization extended beyond the gut. We found a significant increase in the number of B220+ B cells in the bone marrow seven days following the inhibition of GAPs, which was accompanied by the expansion of IgM+IgD-immature B cells without a change in the number of IgM+IgD+ mature B cells (**Fig. 3A-C**). To confirm that the Math1^Cre*PR^ was not expressed in the bone marrow, we generated Math1^Cre*PR^ x ROSA^lsl^ ^dtomato^ inducible reporter mice and evaluated for the presence of dtomato in the bone marrow compartment. We did not detect dtomato expression in any bone marrow cellular population (**Ext. Fig. 3**), indicating that the expansion of B cells was not a bone marrow-intrinsic effect.

**Figure 3:**
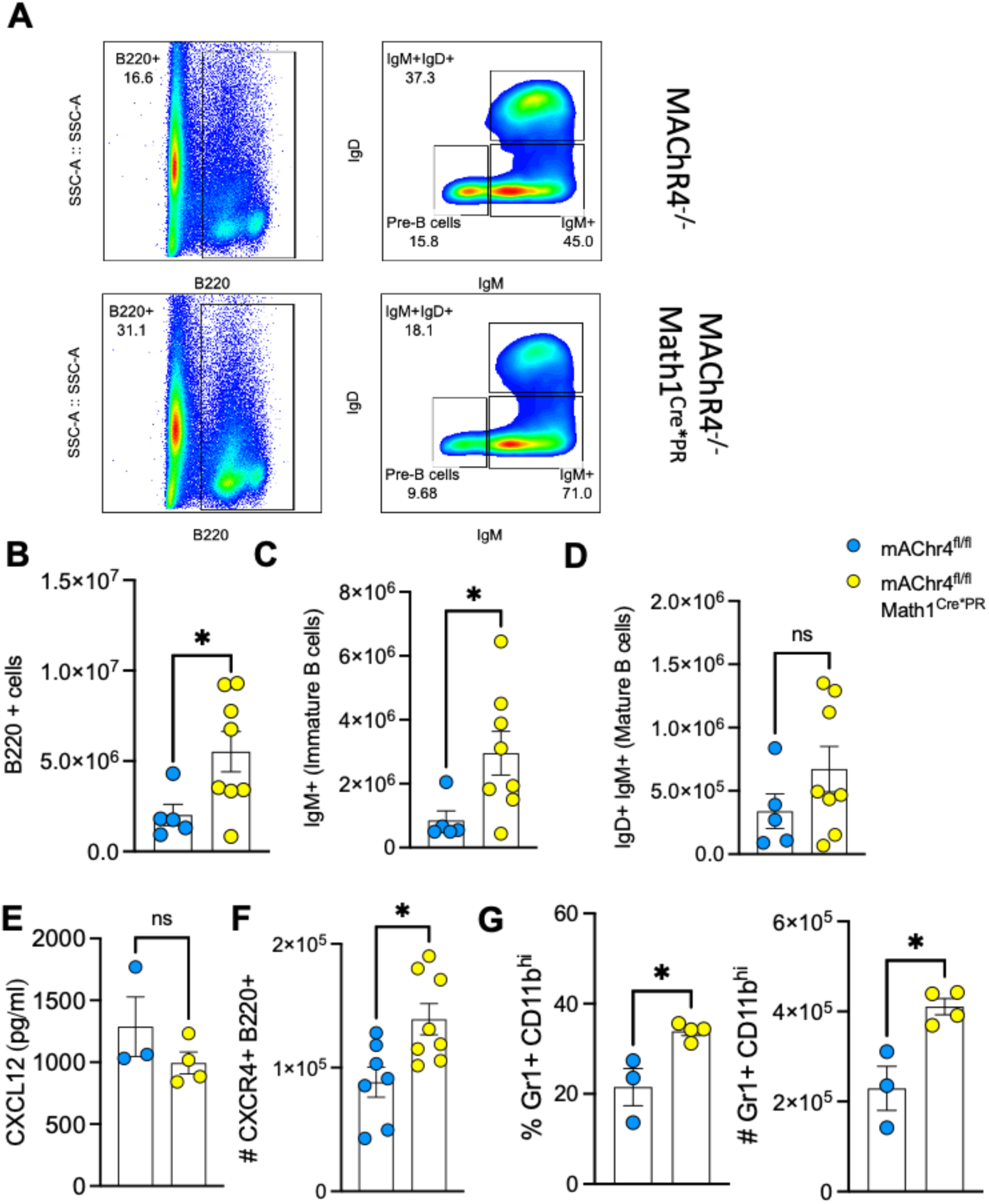
Bone marrow B cell mobilization following alteration of intestinal GAPs. **(A-D)** Bone marrow (B220+), (B220^+^ IgM^+^ IgD^-^, immature B cells) and (B220+ IgM+ IgD+, mature B cells) subsets were quantified following inhibition of GAPs in mAChR4^fl/fl^ Math1^Cre*PR^ mice and littermate controls. **(E)** CXCL12 was measured in the bone marrow extracellular fluid from single femur bone per mouse. **(F)** Expression level of CXCR4 on bone marrow B220+ cells was determined by flow cytometry. **(G)** Frequency and absolute numbers of mature neutrophils (Gr1^+^ CD11b^hi^) were measured in bone marrow of mAChR4^fl/fl^ Math1^Cre*PR^ mice and littermate controls. Data pooled from 2-3 independent experiments. Statistics were calculated by unpaired *t*-test. *p<0.05.

Various chemokines have a critical role in B cell trafficking, both during inflammation and in steady state. The CXCL12-CXCR4 axis is particularly important in controlling B cell homing to and from the bone marrow [34]. CXCL12 is produced by stromal cells and is critical for generation of pre-pro-B-cells and movement of all B cell progenitors within the bone marrow [35]. Hence, we examined the expression of CXCL12 in the bone marrow extracellular fluid following manipulation of intestinal GAPs. While we did not observe any change in the expression of CXCL12, the expression of its receptor CXCR4 was significantly elevated among all the B220+ cells (**Fig. 3E-F).** Given the correlation between emergency granulopoiesis and B cell populations, we examined the frequency of myeloid subsets during the same time frame that we had explored the frequency of B cells. Using the markers Gr-1 and CD11b, we noted significant expansion of mature Gr1+ CD11bhi cells in the bone marrow of mAChR4^f/f^Math1^Cre*PR^ mice compared with littermate controls (**Fig. 3G**). Collectively, these findings show that inhibition of GAPs changes the granulocyte and B cell populations in the bone marrow.

### Plasma cell accumulation in SI-lamina propria is abrogated by FTY720 treatment

Plasma cells accumulate and survive for long periods of time in the SI-Lamina propria and bone marrow and migrate in response to acute injury [33]. To investigate whether plasma cells expanding in the SI upon GAP manipulation are driven by the migration of immune cells from other organs, we treated mAChR4^f/f^Math1^Cre*PR^ and littermate mice with FTY720 following inhibition of GAPs (**Fig. 4A**). FTY720 (fingolimod) is a S1P1 receptor modulator that prevents the migration of leukocytes [36, 37]. We observed that FTY720 abrogated the expansion of plasma cells in the intestine following GAP inhibition (**Fig. 4B**). Additionally, FTY720 abrogated the increased production of IgA by LP cells after GAP inhibition (**Fig. 4C**), consistent with this increase being due to migration of additional IgA-producing plasma cells as opposed to increased production by those already present in the gut. Moreover, FTY720 treatment resulted in the loss of ICOS-production by LP non-Tfh cells in mAChR4^f/f^Math1^Cre*PR^ mice (**Fig. 4D**). Taken together, these studies indicate that the plasma cell expansion we observe upon inhibition of GAPs is dependent upon lymphocyte trafficking from extra-intestinal sites.

**Figure 4:**
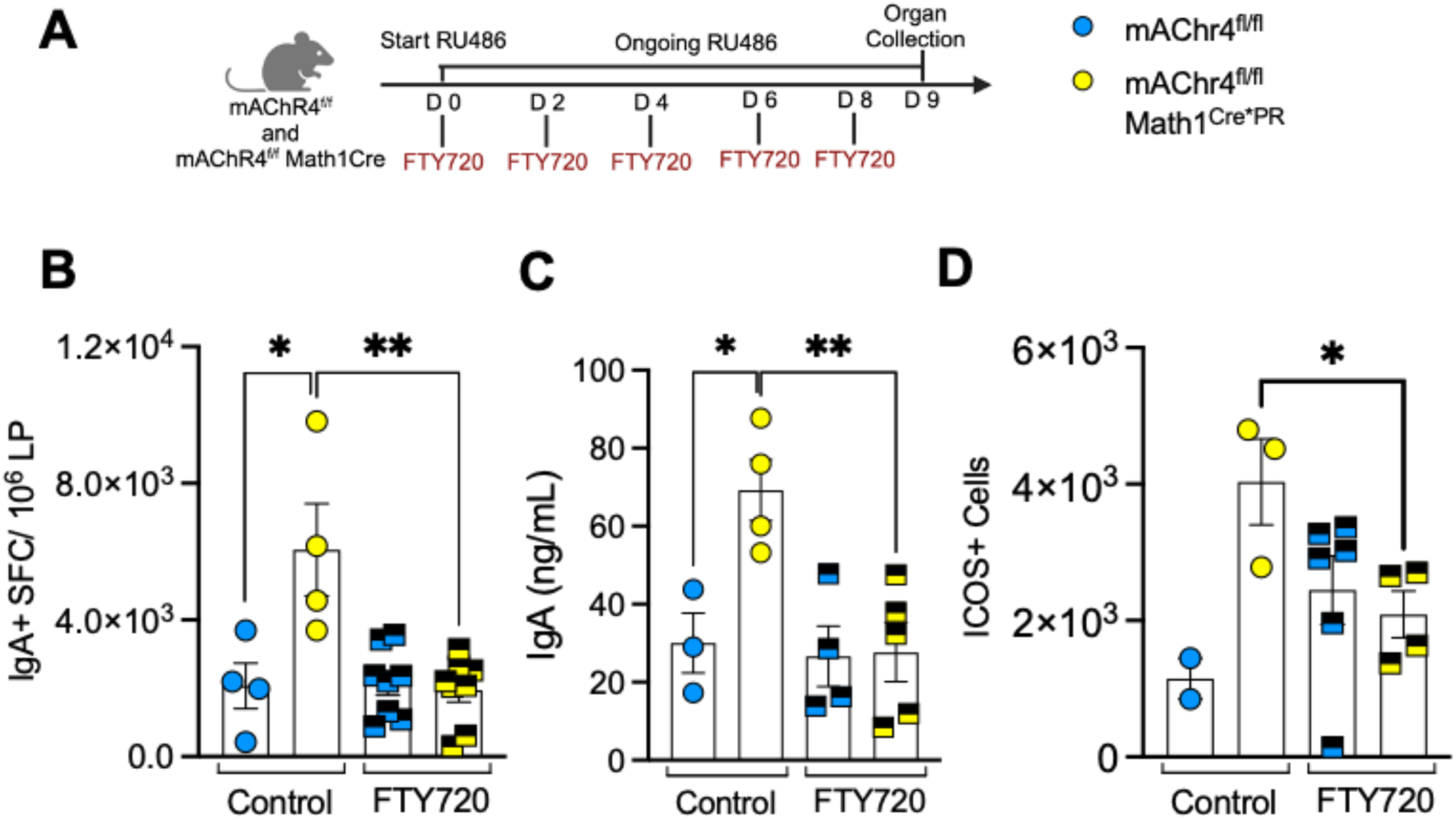
Expansion of plasma cells in the lamina propria is impaired by FTY720 treatment. (A) Schematic overview of experimental setup and animal treatment. (B) Graph depicts small intestinal IgA-secreting cells measured by ELISpot assay in mAChr4^f/f^ and mAChr4^f/f^ Math1^Cre*PR^ mice, treated with PBS or FTY720. (C) ELISA analysis of IgA concentration in small intestinal lamina propria culture for 3 days. (D) quantification of absolute ICOS+ non-Tfh cells. Data combined from 2-3 independent experiments. Statistics were calculated by unpaired *t*-test. *p<0.05, **p<0.01.

### Plasma cell expansion in response to enteric pathogen is driven via goblet cell mediated pathway

We have previously shown that enteric infection with *Salmonella typhimurium* rapidly inhibits small intestinal GAPs in an IL-1β dependent manner [18]. In this study, we observed an expansion of IgA plasma cells in the lamina propria upon *Salmonella* infection (**Fig. 5A**), which was accompanied by increased IgA secretion in the intestinal lumen (**Fig. 5B**). *Salmonella* infection did not influence the levels of circulating or fecal IgA (**Fig. 5 C-D**). In addition, we observed a modest non-significant increase in J-chain levels in the lamina propria cell cultures (**Fig. 5E**). *Salmonella*-infected mice had increased expression of TSLP (**Fig. 5F**), a factor known to promote plasma cells maintenance [38]. In parallel with our earlier observations, we noticed that *Salmonella* infection resulted in increased ICOS production from non-Tfh cells in the small intestine (**Fig. 5G**). Prior studies using *Salmonella* infection models have demonstrated that a portion of marrow resident cells quickly become activated to produce bacterial-specific IgM [39]. Indeed, we observed an expansion of B220+ B cell and specifically IgM+ IgD-immature B cells in the bone marrow, increased CXCR4 expression among B220+ cells (**Fig. 5H-I**) and significantly increased granulopoiesis, as noted by expansion in Gr1+ CD11b^hi^ cells in the bone marrow from infected mice (**Fig. 5J**).

**Figure 5:**
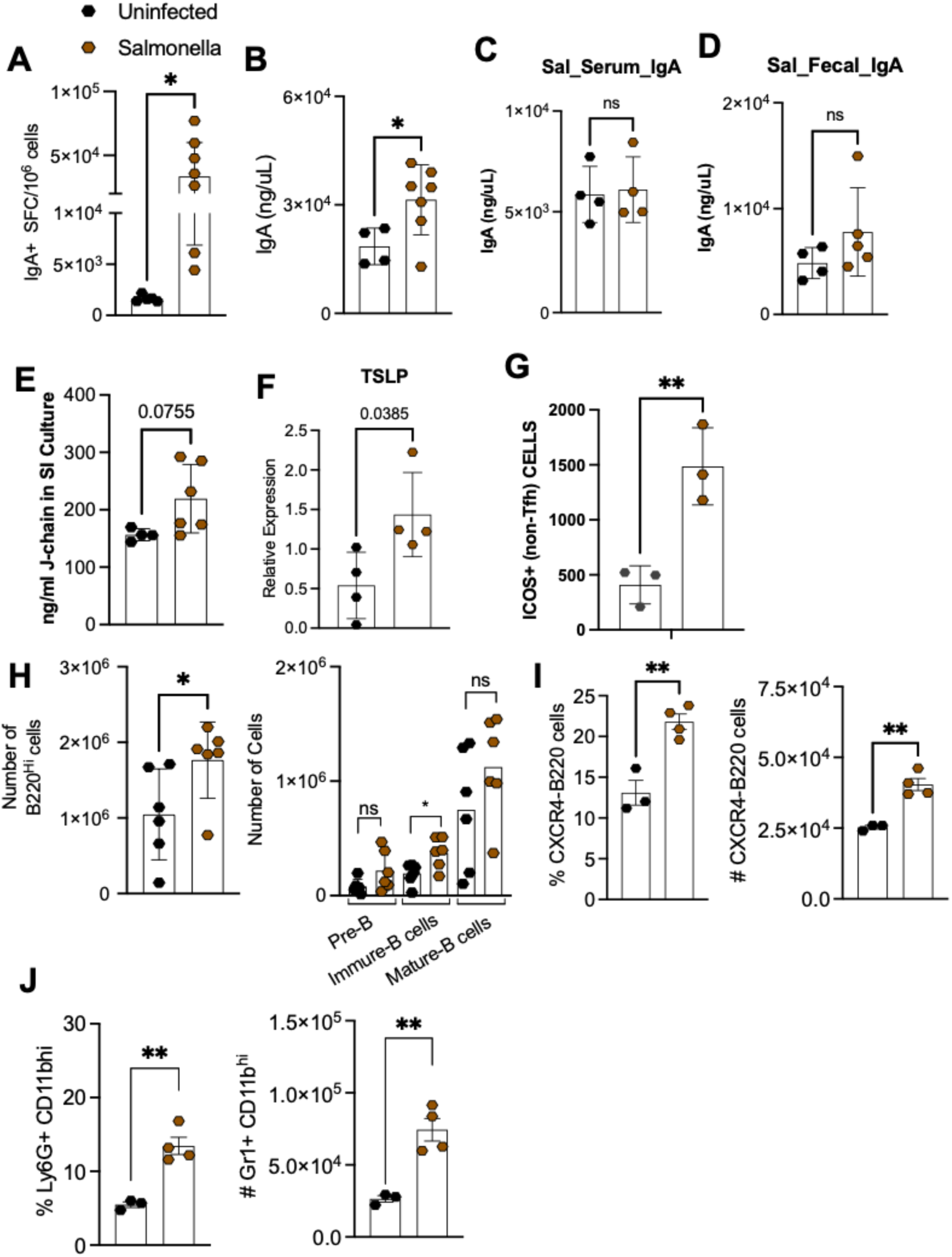
Enteric infection promotes intestinal IgA+ plasma cell accumulation and bone marrow B cell mobilization. **(A)** Graph depicts small intestinal IgA-secreting cells measured by ELISpot assay in C57BL/6 mice infected orally with *Salmonella* for 4 days or in uninfected mice treated with PBS. **(B)** IgA concentration were measured by ELISA in the small intestinal luminal content and in the serum **(C)** of these mice. (**D**) ELISA data from Fecal content of Salmonella infected mice. (**E**) J-chain levels were measured using ELISA in the cultures of small intestinal immune cells from infected and uninfected mice. (**F**) Relative expression of PIGR and TSLP was determined by RT-PCR on small intestinal tissue. (**G**) Flow cytometry analysis of ICOS staining among non-Tfh CD4+ CD45+ cells in Salmonella infection and uninfected mice. (**H**) Bone marrow B220+ cells were assessed following 4 days of *Salmonella* infection. Bone marrow subsets pre-B cell (B220^+^ IgM^-^ IgD^-^), immature B cells (B220^+^ IgM^+^ IgD^-^) and mature B cells (B220+ IgM+ IgD+) were quantified in *Salmonella* infected and uninfected mice. Expression level of CXCR4 on bone marrow B220+ cells was determined by flow cytometry measurement. (**I**) Frequency and absolute numbers of mature neutrophils (Gr1^+^ CD11b^hi^) were measured in bone marrow of Frequency and absolute numbers of mature neutrophils (Gr1^+^ CD11b^hi^) were measured in bone marrow of Salmonella infected and uninfected mice (**J**). Data pooled from 2-3 independent experiments. Statistics were calculated by unpaired *t*-test. *p<0.05, **p<0.01.

Using mice with an inducible deletion of MyD88 on goblet cells (MyD88^f/f^ Math1^Cre*PR^), we have previously shown that if GAP inhibition is overridden during *Salmonella* infection, inflammatory responses are mounted towards dietary antigens, pathogen dissemination is increased, disease course in worsened [18]. Indeed, we observed that treatment with recombinant mouse IL-1β resulted in accumulation of IgA+ plasma cells in the LP of MyD88^f/f^ mice, where GAPs are inhibited by IL-1β, but not in MyD88^f/f^ Math1^Cre*PR^ mice where IL-1β mediated GAP inhibition is prevented (**Fig. 6A**). Therefore, to determine whether GAP inhibition during *Salmonella* infection was contributing in part to the expansion of IgA-secreting plasma cells in the small intestine, we examined plasma cell expansion in the LP of MyD88^f/f^ Math1^Cre*PR^ and littermate control mice following *Salmonella* infection (**Fig. 6B)**. On day 4 after oral infection, the number of IgA-secreting plasma cells were significantly lower in MyD88^f/f^ Math1^Cre*PR^ mice (**Fig. 6C**). This reduction in the number of IgA-secreting plasma cells is accompanied with a trend toward reduction IgA in lamina propria cell cultures (**Fig. 6D).** Thus, inhibition of small intestinal GAPs during *Salmonella* infection is a key part of a physiologic response to infection that promotes the expansion of LP IgA plasma cells.

**Figure 6:**
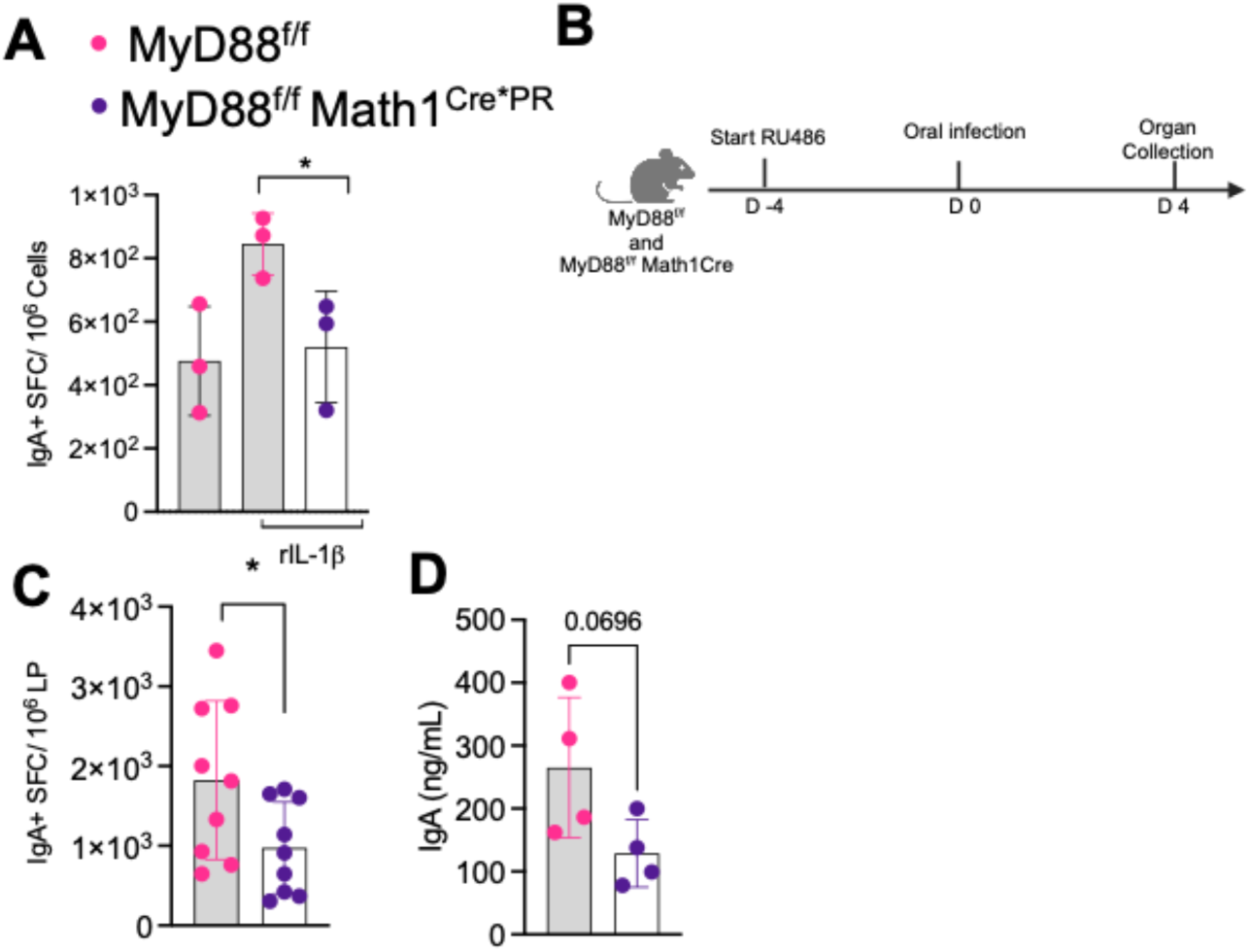
Enteric infection mediated intestinal IgA+ plasma cell accumulation is mediated via intestinal goblet cell signaling. **(A)** Graph depicts small intestinal IgA-secreting cells measured by ELISpot assay in MyD88^f/f^ and MyD88^f/f^ Math1^Cre*PR^ mice treated with rIL-1β for four days or untreated. **(B)** Schematic overview of experimental setup and animal treatment. **(C)** IgA-secreting cells were measured by ELISpot assay MyD88^f/f^ and MyD88^f/f^ Math1^Cre*PR^ mice treated with *Salmonella* for four days. (**D**) Concentration of IgA was measured in small intestinal lamina propria cultured for 3 days. Data pooled from 2 independent experiments. Statistics were calculated by unpaired *t*- test. *p<0.05.

## DISCUSSION

This study reveals an unexpected role for goblet cells and GAPs during enteric infection in promoting humoral immune responses. The findings are unexpected, as to date GAPs have not been observed to have interactions with B cells, and when present, GAPs have not been observed to promote humoral immune responses to luminal antigens. While not explored here, the rapidity of the plasma cell expansion induced by GAP inhibition likely represents mobilization of pre-existing humoral responses as opposed to generation of new responses, although some of the findings seen after GAP inhibition, such as expansion of Tfh cells and ILFs, suggests this cascade of events likely results in enhanced priming of humoral immune responses to new antigens encountered in the gut.

This study extends our understanding of how intestinal epithelial cells respond to an enteric challenge to facilitate immune responses. The gut contains the largest number of immune cells in the body separated by a single layer epithelium from trillions of microbes in the gut lumen. This separation is maintained in part by the gut barrier which consists of the mucus layer, the epithelium, and an immunological barrier. It is interesting that intestinal goblet cells play an integral role in each of these components by secreting mucus, which is a physical component of the single layer epithelium, and by forming and inhibiting GAPs, both of which contribute to adaptive immune responses.

In the steady state, GAPs form continuously in the small intestine due to ACh acting on the mAChR4 of goblet cells and the lack of EGFR activation in goblet cells, which inhibits the ability of goblet cells to respond via the mAChR4 [16]. In contrast, goblet cells in the colon do not form GAPs in the steady state as MyD88-dependent sensing of the microbiota activates EGFR on goblet cells and suppresses responses to ACh via mAChR4 [17]. Notably, GAP inhibition does not impair goblet cell secretion [16], such that the physical and chemical mucus barrier can be maintained during exposure to hostile environments such as enteric infection or the dense microbial load in the colon. In the steady state, ACh does not appear limiting for GAP formation, and control of GAPs largely centers around their inhibition by EGFR activation [17]. The convergence of GAP regulation on the EGFR, which can be activated by multiple stimuli including EGFR ligands, TLR ligands and IL-1R ligands transactivating the EGFR via MyD88 results in the ability to control GAPs in multiple settings to facilitate and coordinate appropriate immune responses [15, 18, 40–43].

The bone marrow is an organ of exceptional immunological importance as it is the site of primary generation of leukocytes as well as harboring long-lived lymphocyte and plasma cell populations with the capacity for rapid mobilization for recall responses. The dynamic nature of the bone marrow enables it to respond in rapid manner with myelopoiesis followed by lymphopoieses in responses to various hostile insults [30]. During sepsis, the protective value of acute antibody production has been shown to be critical for survival [44, 45]. Interestingly, our experiments reveal that B cell production, as well as supportive environmental alterations including expansion in CXCR4 expression and increase in Gr1+ cells (myelopoiesis), occur in response to changes in intestinal goblet cells. Long lived plasma cells generated in the gut can reside in the bone marrow and mobilize upon insult [33]. It is interesting in this context to postulate that inhibition of GAPs may be a unifying event in gut insults that results in the mobilization of humoral immunity in the gut.

We do not understand how signals from intestinal goblet cells are delivered to distant sites such as the bone marrow to mediate these events. Similarly, intestinal epithelial cell endoplasmic reticular (ER) stress was found to induce microbiota-independent expansion of B1 B cells in the peritoneal cavity, culminating in increased IgA production in the LP and secretion in the gut lumen [46], but the mechanism by which these signals are delivered to extraintestinal sites remains elusive. We did not see upregulation of ER stress genes or pathways in our transcriptomic analysis of mice with GAP inhibition. Hence, we do not favor intestinal epithelial ER stress as a component of this cascade. However, future work could uncover that common mechanisms, such as migratory cells and/or cytokines, relay signals to extraintestinal sites in both these processes.

The generation of humoral immune responses in the gut have been areas of intensive investigation, yet how these are controlled, mobilized, and coordinated to respond to insults in the gut is not well understood. Here we describe an unexpected role for intestinal goblet cells in mobilizing humoral immune responses during enteric infection. The role of goblet cell secretion to maintain the mucus barrier has been appreciated for hundreds of years, while only recently has the role of goblet cells to form GAPs been identified. It is nonetheless surprising that this apparently humble epithelial cell has central roles in coordinating adaptive cellular cell responses when forming GAPs and switching to facilitate the mobilization of adaptive humoral responses when GAPs are inhibited, bestowing intestinal goblet cells with important roles in guiding intestinal immunity.

## METHODS

### Animals

All mice were on C57BL/6 background, purchased from The Jackson Laboratory and were bred in house. Mice were housed in a specific pathogen-free facility and fed a routine chow diet. Math1^fl/fl^ mice were bred to vil-Cre-ER^T2^ mice to generate mice with inducible depletion of GCs following deletion of Math1 in villin-expressing cells. Math1^fl/fl^vil-Cre-ER^T2^ mice and the injection protocol to induce GC deletion have been previously described [17]. mAChR4^fl/fl^ mice [47] were a kind gift from Dr. Jurgen Wess (National Institute Health, Bethesda, MD). mAChR4^fl/fl^ mice were bred to Math1^Cre*PR^ mice [48] to generate mice with an inducible deletion of mAChR4 in goblet cells. These mice and Cre-negative littermate controls were orally gavaged with mifepristone (RU486) (10mg/kg) every day starting four days prior to use in experiments. To induce the deletion of MyD88 on intestinal goblet cells, we used a previously published strain of MyD88^f/f^ and MyD88^f/f^ Math1^Cre*PR^ [17, 18]. As indicated above, mice were orally gavaged with mifepristone (RU486).

All mice were bred in house and cohoused littermates were used as experimental controls. Mice were housed in a specific-pathogen-free facility and fed routine chow diet. Animal procedures and protocols were performed in accordance with the IACUC at Washington University School of Medicine.

### Treatments and infection

To assess the effect of FTY720 treatment on plasma cell accumulation and expansion, mice were intra-peritoneally (i.p.) administered with PBS or 100ug FTY720 (Millipore Sigma). At start of the experiment, mice were orally treated with mifepristone (RU486) as described above, with i.p. treatment of PBS or FTY720. Subsequently, PBS or FTY720 dosing was continued every second day till end of the experiment on Day 9.

*Salmonella typhimurium* wildtype strain SB300A1[18, 49] were grown with shaking overnight at 37 °C in Luria-Bertani (LB) broth, sub-cultured for 4 h, and washed twice with cold PBS prior to use. For oral *Salmonella* infections, mice were deprived of food for 4 h and gavaged orally with 5 × 10^7^ CFU of bacteria in 200μl PBS. Mice were kept without food and water for 1 h after infection. The final infection dose was verified by plating serial dilutions of the bacterial suspension on LB agar plates.

To inhibit GAPs using IL-1β treatment, mice were treated i.p. with 100 ng of recombinant murine IL-1β (R&D Systems, Minneapolis, MN). At start of the experiment, mice were orally treated with mifepristone (RU486) as described above for 4 consecutive days. On day zero, MyD88^f/f^ and MyD88^f/f^ Math1^Cre*PR^ mice were treated with IL-1β or PBS, concomitantly with RU486.

### Lamina propria (LP) lymphocyte isolation

LP lymphocyte isolation and intracellular cytokine and transcription factor staining were performed as described previously [15]

Bone marrow (BM) lymphocyte isolation:

Single-cell bone marrow suspensions were made by flushing femurs and tibiae with ice cold PBS. Cells were filtered through a 40 micron cell strainer.

### Flow cytometry

Fluorescently conjugated antibodies were purchased from BioLegend, Becton Dickinson and Invitrogen (eBioscience). Samples were analyzed using an Attune (Thermo Scientific), and data were processed with FlowJo (v10.1).

### Cytokine measurements

Serum was obtained at five-weeks post colonization. To assay cytokine levels in mouse serum, we used multianalyte assay (LEGENDplex Mouse B cell Panel; #740819, BioLegend, San Diego, CA) according to the provided protocol. To collect bone marrow extra-cellular fluid, 1 tibia and femur were cut into small pieces using sharp scissors and resuspended in 100 µl of PBS, followed by centrifuging at 3000rcf. The supernatants were used to measure CXCL12 by enzyme-linked immunosorbent assay (ELISA, R&D Systems).

### ELISpot Assays

ELISpot assays were performed as before. Briefly, ELISpot plates (EMD Millipore MSIPS4W10) were coated with anti-mouse Ig (Southern Biotech). Plates were washed and blocked, before being coated with cells isolated from small intestinal lamina propria. Detection was performed with anti-IgA, IgM and IgG antibodies. ELISpot plates were developed with BCIP/NBT (Sigma, B1911).

### RNA-Sequencing

RNA was isolated from small intestinal tissue (3 cm in length) using the Zymogen RNA isolation kit as per manufacturer instructions. Library preparation and paired end 150bp sequencing was performed by Genewiz. Alignment of RNA sequencing reads on mouse reference genome mm10 was performed using Partek® Flow® software, version 8.0 (Partek Inc., St. Louis, MO). Differential expression analysis was then performed to analyze for differences between conditions and the results were filtered for only those genes with adjusted p-values less than or equal to 0.05.

### Quantitative RT-PCR

mRNA extraction was performed using a RNeasy Plus Mini Kit (Qiagen, Germantown, MD), and complementary DNA (cDNA) synthesis was performed using the High-Capacity cDNA reverse transcription kit (Thermo Fisher Scientific) according to the manufacturer’s recommendations. Each cDNA sample was diluted by fivefold after cDNA synthesis, and real-time PCR of each gene was carried out in duplicate on a QuantStudio 3 (Thermo Fisher Scientific) using SYBR Green (Power SYBR Green, Applied Biosystems). Ct values were read and exported from the QuantStudio software, and the commonly used 2(-ΔΔ*Ct*) method was used to calculate relative gene expression using *Gapdh* or *Hprt1* as the housekeeping gene. Primer sets utilized for real time PCR are listed below:

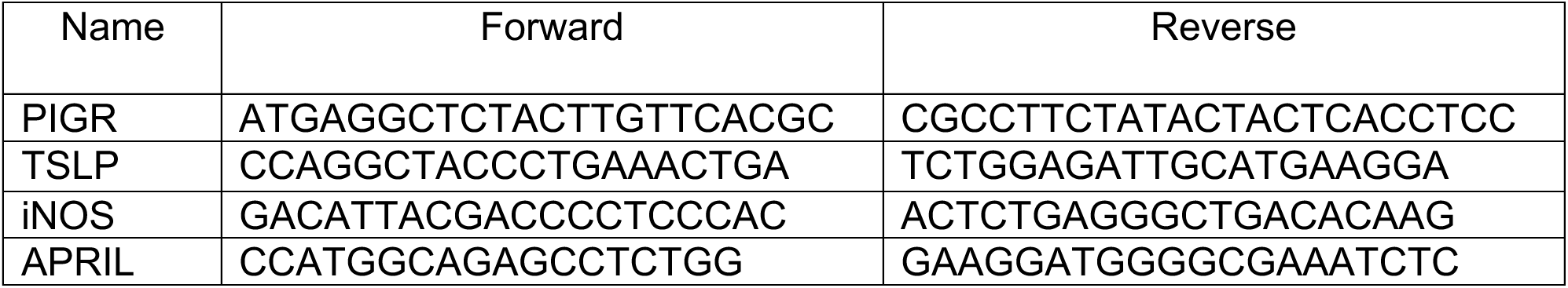

### ELISA

Serum, fecal or small intestinal-luminal content, or lamina propria cells cultured for 3 days were used to detect IgA, IgM or IgG antibody levels. ELISA was conducted as described previously[22]. In short, 96-well Immuno 4 plates (Fisher Scientific) were coated with 10mg/ml goat anti-mouse Ig (Southern Biotechnology) and blocked with PBS containing 5% BSA. The above mentioned samples were diluted in PBS containing 1% BSA and 0.05% Tween 20 and incubated for 2 hours at room temperature. Following incubation with detection antibody, plates were developed using p-nitrophenyl phosphate alkaline phosphatase substrate (Sigma-Aldrich). Each sample was measured in duplicate in two - three dilutions, and two-three independent experiments were performed.

### ILFs

Small intestinal tissue ILFs were measured as described before [22, 50].

### Extraction of fecal genomic DNA and profiling the fecal microbiota

Luminal contents were collected and processed for DNA isolation using the Power Fecal DNA Kit (Qiagen). The V4 region of 16S rRNA gene was PCR-amplified using barcoded primer described previously and sequenced using the Illumina Miseq Platform (2 x 250-bp paired-end reads). Metagenomic analysis was performed via Kraken Programs performed using Partek® Flow® software, version 8.0 (Partek Inc., St. Louis, MO).

### Statistical Analysis and Data Availability

Data are presented as scatterplots showing individual data points, and dispersion is shown by means ± SD. Unpaired *t* test was used for comparing two groups, assuming equal variance, and ordinary one-way analysis of variance (ANOVA) test with multiple comparisons testing was used for more than two groups. A *P* value of <0.05 was considered significant. Prism v9.3.0 (GraphPad, San Diego, CA) was used for statistical analysis.

The microbiota 16S rRNA sequencing data will be submitted to NCBI Short Read Archives (SRA), which will be publicly available. The RNA-seq dataset will be deposited and publicly available on NCBI GEO.

### Study Approval

Animal procedures and protocols were performed in accordance with the IACUC at Washington University School of Medicine.

## Acknowledgements

This work was funded by grants NIDDK K01-DK133670, Crohn’s and Colitis Foundation Career Development Award 610605 (to D.H.K); Crohn’s and Colitis Foundation Research Fellowship Award 902790 (to V.J); grants R01DK097317, U01AI163073, R37AI112626, and R01AI173220 (to R. D. N).

## Author contributions

DHK performed immune cell phenotyping, and data analysis. DHK, KT, AF, EJ, HK, RM, VV, VJ, BB, SU and DH performed animal models. KT and DHK performed and analyzed the 16S rRNA analysis. KGM did ILFs assessment. DHK performed RNAseq analysis. DHK and RDN designed the experiments, analyzed, and interpreted the data, and wrote the manuscript.

## Notes

### Competing Interest Statement

The authors have declared no competing interest.

