## Supplement for "Small Intestinal Goblet Cells Control Humoral Immune Responses and Mobilization During Enteric Infection"

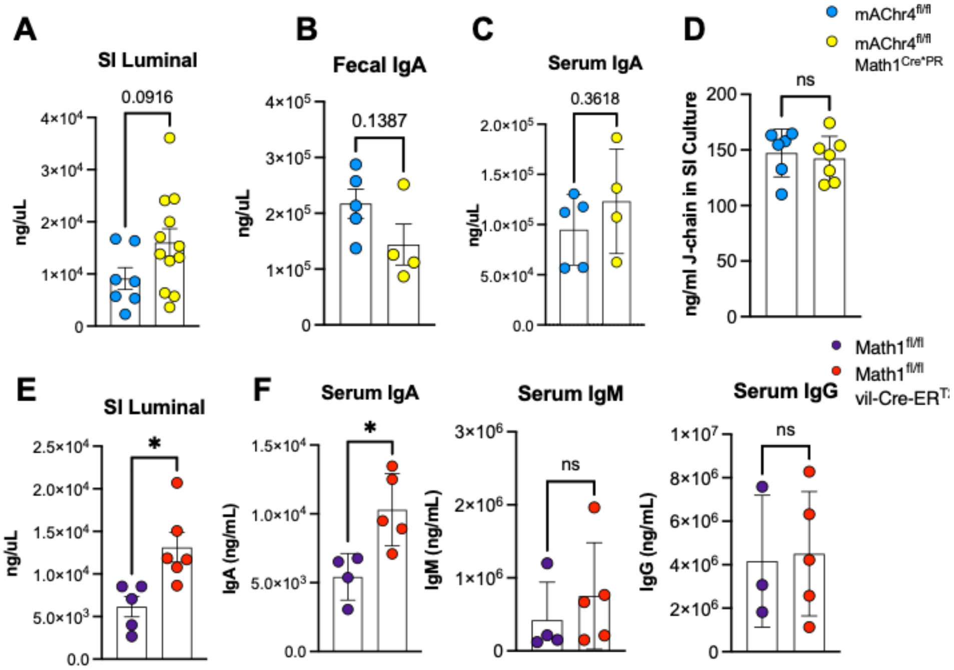


**Extended Figure 1: Effect of goblet cell deletion or inhibition of GAPs on concentration of luminal and systemic IgA. (A-C)** Concentration of IgA based off ELISA assay in (**A**) SI luminal content, **(B)** fecal pellet and **(C)** serum, were measured in mAChr4$^{f/f}$ and mAChr4$^{f/f}$ Math1$^{Cre*PR}$ mice treated with RU486 for seven days. (**D**) J-chain levels were measured using ELISA in the culture supernatants of small intestinal immune cells. (**E**) Concentration of IgA in the small intestinal luminal content and (**F**) IgA, IgM and IgG based off ELISA assay in serum from Math1$^{f/f}$ and Math1$^{f/f}$ Vil-Cre-ER$^{T2}$ mice treated with tamoxifen to induce goblet cell deletion. Statistics were calculated by unpaired *t*-test. *p<0.05.

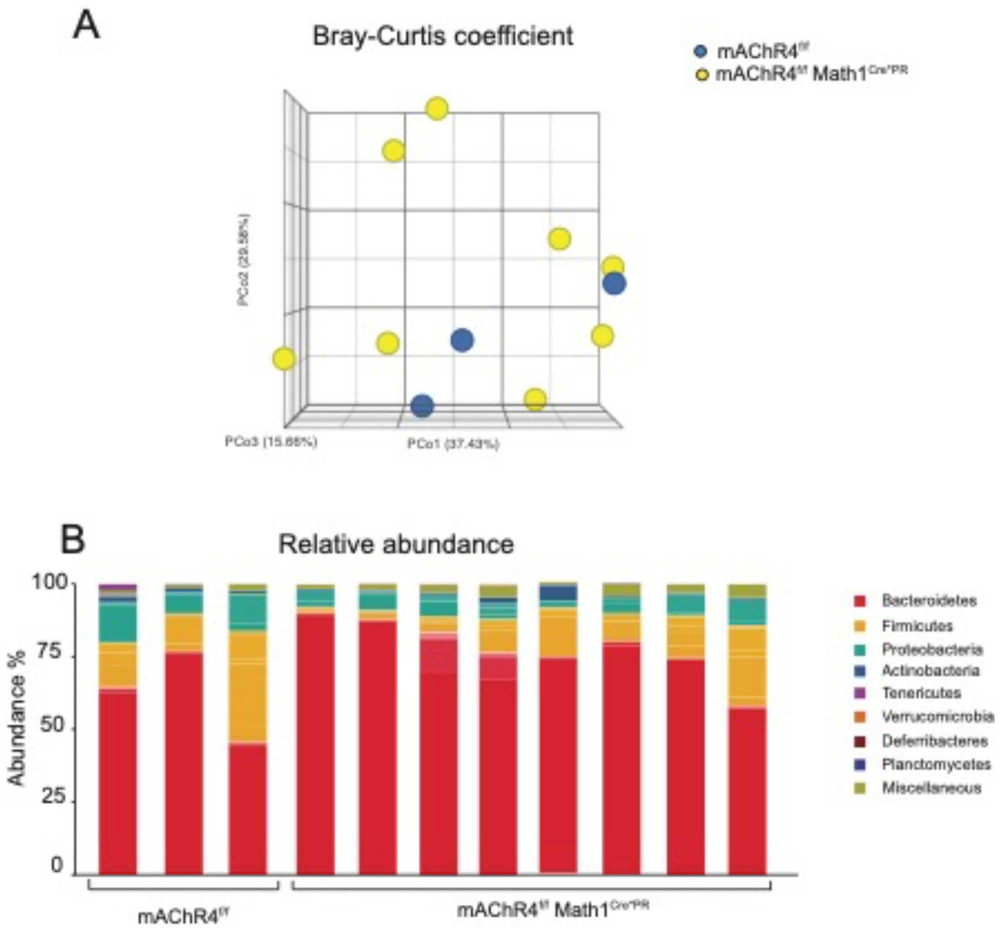


**Extended Figure 2: Inhibition of GAPs does not alter intestinal microbial composition. (A)** Principle Coordinate Analysis using the Bray-Curtis dissimilarity coefficient showing microbial clustering of ileal luminal microbiome, and **(B)** relative abundance of different phyla in the ileal luminal microbiome from mAChr4$^{f/f}$ and mAChr4$^{f/f}$ Math1$^{Cre*PR}$ mice treated with RU486 to activate Cre for seven days.

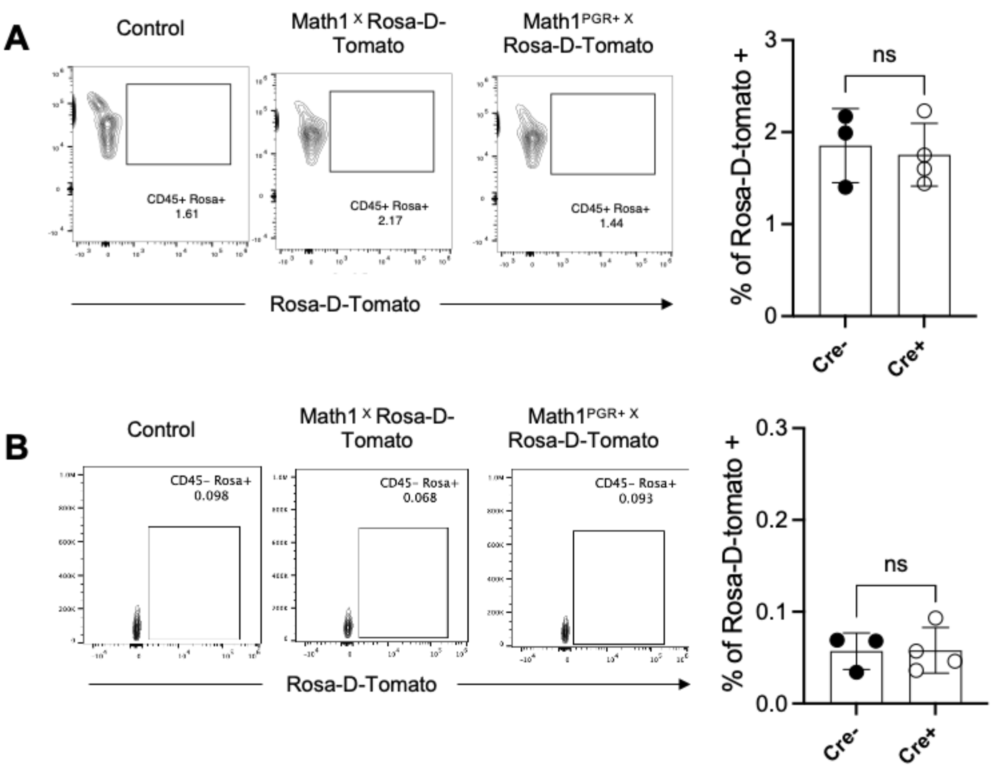


**Extended Figure 3: Bone marrow cells do not express Math1.**

**(A)** Flow cytometric analysis of the of bone marrow cells for surface expression of Rosa-d-tomato among CD45+ cells and in **(B)** among the CD45 negative cell subset in Math1X Rosa-DTomato and Math1CreX Rosa-Dtomato. Statistics were calculated by unpaired *t*-test.
